# Rapid microfluidic perfusion system enables controlling dynamics of intracellular pH regulated by Na^+^/H^+^ exchanger NHE1

**DOI:** 10.1101/2024.10.18.619062

**Authors:** Quang D. Tran, Yann Bouret, Xavier Noblin, Gisèle Jarretou, Laurent Counillon, Mallorie Poët, Céline Cohen

## Abstract

pH regulation of eukaryotic cells is of crucial importance and influences different mechanisms including chemical kinetics, buffer effects, metabolic activity, membrane transport and cell shape parameters. In this study, we develop a microfluidic system to rapidly and precisely control a continuous flow of ionic chemical species to acutely challenge the intracellular pH regulation mechanisms and confront predictive models. We monitor the intracellular pH dynamics in real-time using pH-sensitive fluorescence imaging and establish a robust mathematical tool to translate the fluorescence signals to pH values. By varying flow rate across the cells and duration for rinsing process, we manage to tweak the dynamics of intracellular pH from a smooth recovery to either an overshooting state, where the pH goes excitedly to a maximum value before decreasing to a plateau, or an undershooting state where the pH is unable to recover to ∼ 7. We believe our findings will provide more insight into intracellular regulatory mechanisms and promote the possibility of exploring cellular behavior in the presence of strong gradients or fast changes in homogeneous conditions.

## 1 Introduction

Biological processes are nearly all pH dependent, and hence, this is why this parameter is so tightly controlled. ^1,2^ Quantitatively, pH represents the abundance of the smallest cation in the universe in aqueous solution and is measured as the cologarithm of free H^+^ concentration. At the cellular level, pH is both one of the most finely tuned parameters and one of the most challenging to control. Indeed, (i) many reactions result in the rapid release or consumption of protons and/or acidbase equivalents, (ii) respiratory complexes use H^+^ gradients across membranes to produce and store energy and (iii) pH determines the protonation state of macromolecules and thereby dictates their surface charge that is key for their interactions. Finally, pH is also involved in the catalytic activity of enzymes that use acid-base catalytic mechanisms and/or is a critical allosteric regulator of enzymes or membrane proteins. There are numerous reviews on pH regulation, which provide invaluable insights in both normal and pathological conditions. ^3–5^

To achieve the challenge of finely tuning pH evolution in time and space, evolution has selected a large group of proteins capable of transporting acid-base equivalents across membranes. These include pumps, ^6^ leaks and secondary transporters ^7–9^ that can translocate H^+^, bicarbonate ^10^ or others buffers. In this context, we have previously built a fully tractable mathematical model for pH regulation that in particular predicts overshoots of pH recovery when a transient change that produces an acidification is suddenly removed. ^11^ Such and important property is biologically important as it could lead to pathological situations if not adequately damped, possibly by functional redundancy.

To challenge such a mechanism in detail, it is crucial to measure intracellular pH at steady state but even more so to be able to measure the dynamics of pH change depending on cellular conditions. The main gold standard technique to get reliable quantitative pH measurements is based on the use of pH sensitive fluorescent probes, whose absorption and/or emission spectra depend on their protonation state. ^12^ Different measurement methods and devices are based on such pH-dependent spectral properties of probes, for e.g. fluorometers or fluorescence-equipped microplate readers, but one of the most widely used, due to its ability to image single cells, is based on live fluorescence videomicroscopy equipped with high-resolution camera and ad hoc illumination systems. ^13^ Although videomicroscopy is widely used, it has certain limitations, particularly when measuring pH changes triggered by very transient disturbances, which therefore require rapid kinetics. Paradoxically, the limits do not presently reside in cameras or illumination systems but in the ability to use very rapid perfusion systems that allow extremely fast exposure to solutions and in parallel the elimination of the incubation medium at the same rate.

The work presented here allows for the first time to access this valuable information on rapid intracellular pH variations that enables to document the conditions leading to the occurrence of overshoots. Intracellular pH overshooting is detrimental in the maintenance of cellular homeostasis and functionality, as it leads to a transient alkalinization that follows the acidification, both of them being detrimental, as they do not correspond to a physiological pH. We previously showed that such overshooting is a more general property of biological regulation mechanisms. Measuring rapid pH overshooting provides a unique window into the understanding of these phenomena that are extremely important and widespread at all scales in living systems. ^2,11^ Actually, the homeostasis of the cell arises from the steady state of competing regulatory phenomenons that are membrane transporters and chemical reactions. In a previous work, ^11^ we proved that alteration of this physiological regulation by a vanishing perturbation - meaning that we start and finish to the exact same experimental parameters - yields a pH overshoot during its relaxation. The dimensions of these overshoots depend on the characteristics of the perturbation that we impose, but neither their amplitude nor their time occurrence obey an analytical expression.

Microfluidics has been a powerful tool allowing for on-chip cell culture and real-time monitoring of extracellular pH. ^14,15^ Taking its advantages, in this study, we developed a laminar flow regime using microfluidics to impose precise and reproducible adequate perturbations in order to quantify their effects on intracellular pH. We performed a 3-step regulation of intracellular pH in a NHE forward activity (Fig. 1), in which protons were extruded from the cytosol to the extracellular environment via a Na^+^/H^+^ exchanger (NHE1). Our system allowed for rapid perfusion of flow through on-chip cultured cells, featuring real-time monitoring of the intracellular pH. We induced various dynamics of intracellular pH by varying flow rates across the cells or duration for rinsing process, including producing an overshooting state that is challenging to obtain in a traditional perfusion system.

**Fig. 1.**
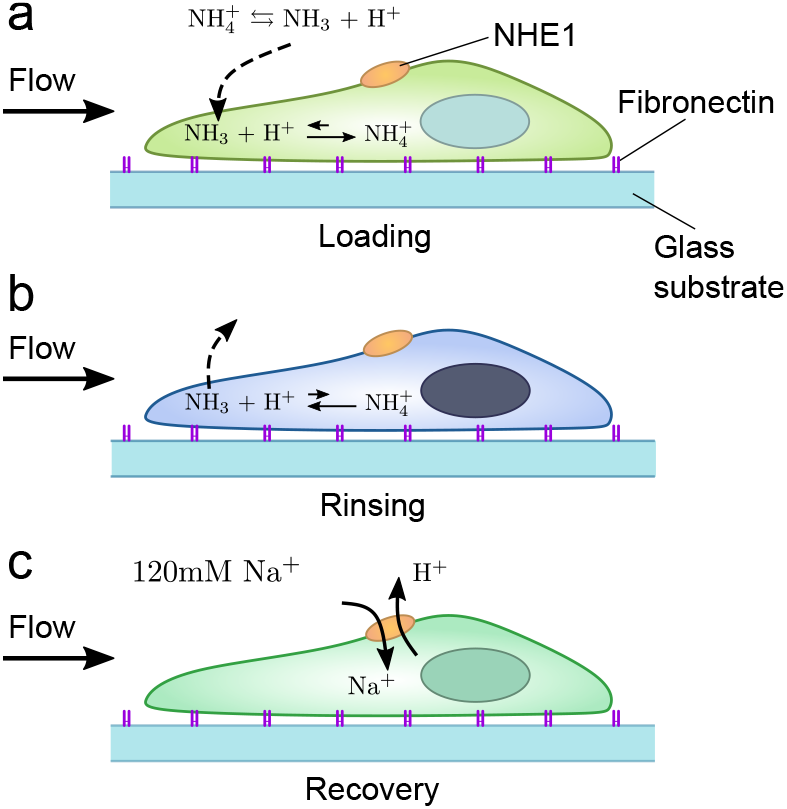
Schematics of the intracellular pH regulation in 3-step NHE forward activity: a) Loading of 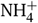 which results in NH_3_ entering through cell membrane and absorbing intracellular protons; b) rinsing of NH_3_ to release protons inside intracellular region; c) recovery of intracellular pH by exchange of supplied Na^+^ ions for intracellular protons through NHE1.

## 2 Methods and Materials

### 2.1 Microfluidic design and fabrication

We designed a microfluidic device to feature on-chip cell culture and live imaging (Fig. 2(a)). The device involves a straight microchannel with height of 50 µm, which is separated into 3 regions of different width. Two large regions with 500-µm width for cell culture and observation purposes are placed at two ends of a thin channel (50-µm width) to enhance the hydrodynamic resistance of the channel. The high resistance helps maintain a slow flow of culture medium across the whole channel when there is a slight pressure difference between the inlet and outlet. We selected the single-channel design over the one with multiple inlets as we need to completely remove the dead volume and prevent the solution mixing or diffusion during the pH regulation experiment.

**Fig. 2.**
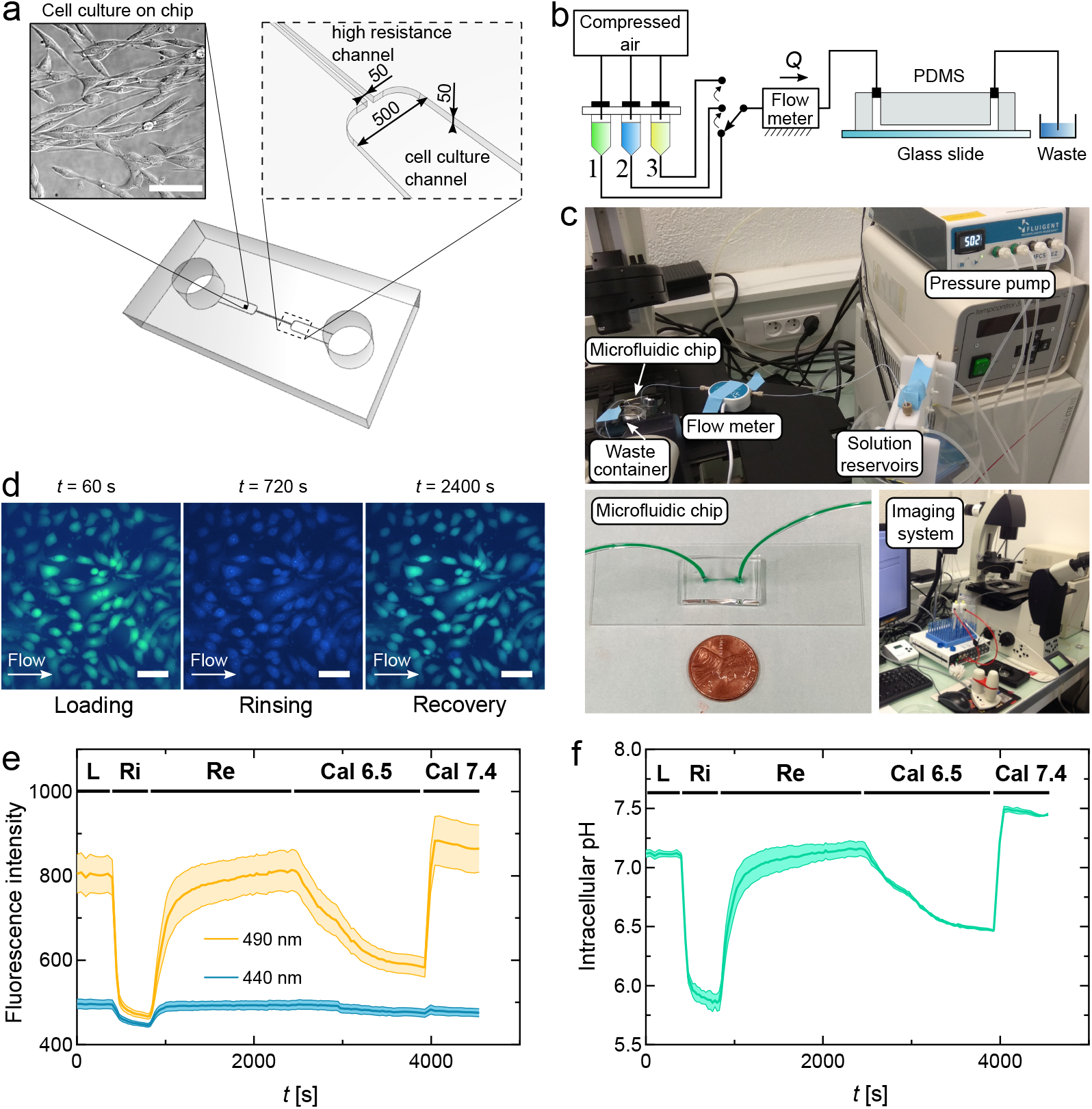
Experimental setup for intracellular pH measurement. (a) Design of the microfluidic device, which consists of a straight channel with height of 50 µm and 2 cell culture regions (*w* = 500 µm) and a high resistance region (*w* = 50 µm) in between. Cells were cultured on a chip for 2 days. Channel dimensions are in µm. (b) Schematic of the microfluidic setup. Continuous flow was applied to eliminate all dead volumes and unwanted mixtures of solutions. (c) Images of the actual overall setup (top), the physical microfluidic chip (bottom left) and the imaging system (bottom right). (d) BCECF fluorescent dye was used to quantify the pH values of cells. BCECF-stained cells change color according to their pH levels at different pH regulation activities. Scale bar: 50 µm. (e) Fluorescence intensity of representative cells from (d) was recorded simultaneously in two wavelengths 490 nm and 440 nm throughout the whole experiment including the Loading, Rinsing, Recovery processes and two calibration processes at pH 6.5 and pH 7.4. (f) Translated intracellular pH values of the representative cells during the whole pH regulation experiment using equation 1. Thick solid lines represent means, and filled areas between thin solid lines represent standard deviations over 8 representative cells. Abbreviations: L – Loading, Ri – Rinsing, Re – Recovery, Cal 6.5 – Calibration pH 6.5, Cal 7.4 – Calibration pH 7.4.

Our microfluidic device was fabricated in PDMS (Sylgard 184, Dow Chemical, Midland, MI, USA) following the standard soft-lithography technique. ^16–18^ A silicone mold was first obtained by photo-lithography with SU-8 100 (MicroChem, Newton, MA, USA) to deposit the design pattern on a silicon wafer with a desired height of 50 µm. We then produced PDMS chips by mixing PDMS monomers with the curing agent at ratio 10:1 and pouring onto the mold. Being cured after incubation at 70 °C for 2 hours, the PDMS chip was then punched with holes at the inlet and outlet. We sterilized the PDMS chip together with a glass slide by autoclave and then bonded them together using a plasma cleaner (Harrick plasma).

### 2.2 Surface treatment for on-chip cell culture

Right after the plasma bonding, a fibronectin solution (F1411, Sigma-Aldrich, St. Louis, MO, USA) diluted at 0.1 mgmL^*−*1^ in distilled water was injected into the microfluidic channel. The device was then incubated overnight at room temperature. The fibronectin coating enhances cell adhesion to the device substrate, thus securing cell positions under flow. The coated device needs to be rinsed with PBS and then filled with culture medium in order to be prepared for cell seeding.

### 2.3 Procedure and reagents for the 3-step NHE forward activity

To facilitate the pH regulation assays for NHE forward activity (Fig. 1), we prepared 3 solutions according to 3 steps: (1) Perfusion of 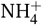 loading solution (Loading solution in short) consisting of 50 mM NH_4_Cl, 70 mM choline chloride, 5 mM KCl, 1 mM MgCl_2_, 2 mM CaCl_2_, 5 mM glucose and 15 mM MOPS at pH 7.4. This step results in NH_3_ entering cells through the membrane and absorbing H^+^ to form 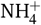. (2) Perfusion of Rinsing solution (120 mM choline chloride, 5 mM KCl, 1 mM MgCl_2_, 2 mM CaCl_2_, 5 mM glucose and 15 mM HEPES at pH 7.0), which allow NH_3_ to escape and release H^+^ inside cells. (3) Perfusion of Recovery solution (120 mM NaCl, 5 mM KCl, 1 mM MgCl_2_, 2 mM CaCl_2_, 5 mM glucose and 15 mM HEPES at pH 7.4), which supplies Na^+^ to the system and establishes the H^+^ extrusion to extracellular environment via NHE1. At the end of each experiment, we applied two pH calibration solutions consisting of 140 mM KCl, 20 mM HEPES, and 5 µM nigericin with pH values adjusted at 6.47 and 7.44. Nigericin helps adjust intracellular pH by exchanging K^+^ and H^+^ ions across cell membranes via passive diffusion.

### 2.4 Cell culture, cell suicide and on-chip seeding

In this study, we used PS120 fibroblast cells stably expressing mutant NHE1, a transfected cell line that has been used in our previous studies. ^11,19,20^ Cells were first cultured in DMEM enriched with 7.5% fetal calf serum and 50 µgmL^*−*1^ PenStrep at 37 °C and 5% CO_2_. Transfection of NHE1 was performed using Lipofectamin 2000 following the protocol from the manufacturer (ThermoFisher, Waltham, MA, USA).

We performed a cell suicide process, which has been described by Milosavljevic et al. ^13^ to select and amplify the population of stable NHE1-expressed cells. To summarize, we first incubated the cells for 1 hour in Loading solution at 37 °C inside an incubator without CO_2_ supply. Next, we removed the Loading solution and washed the cells twice in the Rinse solution. Finally, we replaced the Rinse solution with Recovery solution and incubated for another 1 hour also without CO_2_ supply. CO_2_ was not allowed in this process due to the acidification resulting from the interaction between CO_2_ and the buffers. The Recovery solution was then aspirated and replaced by fresh culture medium so that the cells continued growing. We performed two cycles of this selection process per week. Cells that do not express a functional NHE1 cannot recover a neutral pH level, thus becoming fatal. Cells that survive the acidification through this two-cycle selection express fully and stably the NHE1 protein and function. Experiments in biochemistry, protein characterization, pharmacology to further confirm the proper functionality and expression of NHEs have been conducted over the past 4 decades. ^7^

Upon cofluent, the cells were trypsinized and suspended in culture medium, then seeded into the microfluidic channel by pipetting. We incubated the microfluidic device containing the cells at 37 °C and 5% CO_2_ for 30 minutes. A droplet of fresh culture medium was placed at the inlet of the microfluidic channel, which helped create a continuous flow of fresh medium across the channel and promote cell growth. The high resistance channel is important here as it prevents the droplet from being rapidly depleted and maintains the slow flow of medium for several days. We cultured the cells inside the microfluidic device for 2 days and refilled the droplet every day.

### 2.5 Fluorescence pH imaging

To measure the intracellular pH (pH_i_), we utilized a ratiometric pH-sensitive fluorescence dye BCECF/AM (B1170, ThermoFisher), which has double excitation and is permeable to cells. The pH measurement was conducted by obtaining the ratio between the emission intensity of the dye when excited at 490 nm and at its isosbestic point of ∼ 440 nm. This method normalizes the variation of fluorescence intensity among cells since only the fluorescence ratios are directly translated into pH values. We stained the cells grown inside the microfluidic device by pipetting 200 µL of BCECF/AM diluted in Loading solution at final concentration of 5 µLmL^*−*1^ to the channel inlet. The device was then placed inside an incubator at 37 °C without CO_2_ supply for 30 minutes. Before each experiment, we rinsed the microchannel with the Loading solution to remove the excess dye.

### 2.6 Experimental setup

The microfluidic device containing BCECF-stained cells was connected to our microfluidic system as shown in Fig. 2(a, b). The input flow was applied by a Fluigent pressure control pump (Fluigent, Le Kremlin-Bicêtre, France) integrated with a flow meter, which is able to precisely generate a desired flow rate ranging from 2 to 12 µLmin^*−*1^. The pressure pump and flow meter were connected to a computer and controlled by MAESFLO software. During experiment, solutions were able to be changed quickly by switching the tubing connectors from one solution to another. The output was connected to an open waste reservoir.

Fluorescence pH imaging of the cells was conducted by an inverted microscope (LEICA DMI6000 B) equipped with 440 nm and 490 nm narrow band interference filters and captured by a high-sensitivity camera (Hamamatsu ORCA-Flash4.0 LT+). Fluorescent cell images and the ratio between emission intensity between 490 nm and 440 nm wavelengths were recorded in real-time using the Leica Application Suite software. The pH regulation was performed by applying a single constant flow rate *Q* of Loading, Rinsing and Recovery solution across the microfluidic channel, respectively. We started with the Loading process for 10 minutes, then switched to Rinsing for a duration *t*_r_, and finally switch to Recovery. We recorded the data until the fluorescence intensity was stabilized after Recovery. Fig. 2(c) shows images of cells changing their color according to different processes of the pH regulation experiment. At the end of the experiment, we flowed 2 calibration solutions containing nigericin at pH 6.5 and 7.4 into the channel, respectively. We illustrated the acquired data of each experiment in Fig. 2(d).

## 3 Results and discussion

### 3.1 Establishment of a rapid microfluidic perfusion system for intracellular pH measurement

We achieved a lab-on-a-chip system for real-time pH_i_ regulation assays, including cell culture directly inside the device, high precision controllable flow application with different solutions across the cell chamber, and live imaging feature with fluorescence microscopy and versatile data acquisition on multiple cells (Fig. 2). With microfluidics, we were able to fully control the flow rate and exposure time for each pH regulation solution to interact with the cells, which was not possible for previous traditional perfusion systems. ^13^ Moreover, microfluidic perfusion prevents risks of cross-contaminating solutions, disrupting fluorescence signal reliability and fluctuating sample positions, which usually happens in traditional pipetting methods. By staining the cells with double excitation pH-sensitive fluorescence dye BCECF/AM and using fluorescence microscopy, we captured the cells changing their fluorescence intensity at two different wavelengths (440 and 490 nm) during pH regulation. The software used in the microscopy system (Leica) made it possible to easily select multiple Regions of Interest (ROIs) to record and display data of fluorescence intensity of the two wavelengths in real-time. ^13^ This helped us observe the behavior of multiple cells and control the flow application according to various experimental purposes.

### 3.2 Estimation of intracellular pH values directly from fluorescence intensity data

From the ratio of emission intensity between the two wavelengths obtained from the live imaging, the pH_i_ data of each individual ROI are calculated using the following equation:

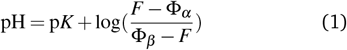

where p*K*≈ 7 for BCECF/AM, *F* is the emission intensity ratio measured at each ROI, Φ_*α*_ is the intrinsic most acidic ratio of the dye, and Φ_*β*_ is the intrinsic most basic ratio of the dye. In this estimation, we need to obtain Φ_*α*_ and Φ_*β*_, which requires two ratio values *F*_*a*_ and *F*_*b*_ at two calibration points, one under acidic conditions (*h* = *h*_*a*_), one under basic conditions (*h* = *h*_*b*_) with *h* = [H^+^], and both points are far from the experimental zone. With *F*_*a*_ and *F*_*b*_ obtained from experiment, we derive Φ_*α*_ and Φ_*β*_ from the following equations:

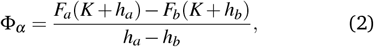

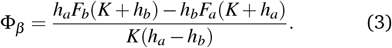

Theoretically, there are no values of ratios *F* that exceed Φ_*α*_ or Φ_*β*_, we can apply equation (1) for the hold set of experimental ratios *F* and the pH value can always be defined (Fig. 2(e)). See Supp. Materials for more details of the calculation.

### 3.3 Different dynamics of intracellular pH observed during the recovery process

We studied the behavior of pH_i_ of randomly selected cells under different flow rates generated by the pressure pump and flow meter. During the 10 minutes of the Loading process, the pH_i_ remained stable, then started to drop when the Rinsing solution reached the cells. We first selected the rinsing duration *t*_r_ = 6 min so that the decreasing pH had enough time to reach a steady state. The Recovery solution was applied at a constant flow rate right after the rinsing duration finished. The Na^+^ ions added by perfusion of the Recovery solution initiated the increase of pH_i_, demonstrating the ion exchange via NHE1 at the cell membrane. ^7,21^ Interestingly, under different flow rates, the pH_i_ recovered with different behaviors (Fig. 3(a)). First, we observed that at *Q* = 4 to 8 µLmin^*−*1^, the pH_i_ gradually increased to a plateau around 7, which is similar to its common behavior observed in traditional perfusion systems. ^13^ We name this behavioral state of the pH_i_ as “Normal recovery.” Secondly, for higher *Q* (≥8µLmin^*−*1^), the recovery was modified: the pH increased drastically to a maximum value, which exceeds the intended pH value before decaying and approaching a plateau (Fig. 3(a) and Supp. Movie 1). This pH regulation is called “Overshooting” phenomenon of the pH_i_. This result confirms the theoretical model introduced by Bouret et al. ^11^ predicting pH_i_ can perform overshooting in regulatory loops. However, the overshooting requires a relatively high-speed continuous flow, which might be the reason for this phenomenon to be challenging to obtain in traditional perfusion systems, which apply flow by gravity and do not remove the incubation medium at enough speed that results in some uncontrolled mixing. Finally, when *Q* was very low (≤ 3µLmin^*−*1^), the pH_i_ recovered very slowly and was not able to reach pH 7, namely “Undershooting” recovery. In many cases of undershooting, the poor pH_i_ recovery often resulted in cell death. Overall, using microfluidics we have unraveled robustly multiple behavioral states of pH_i_ recovery in cells under different flow conditions.

**Fig. 3.**
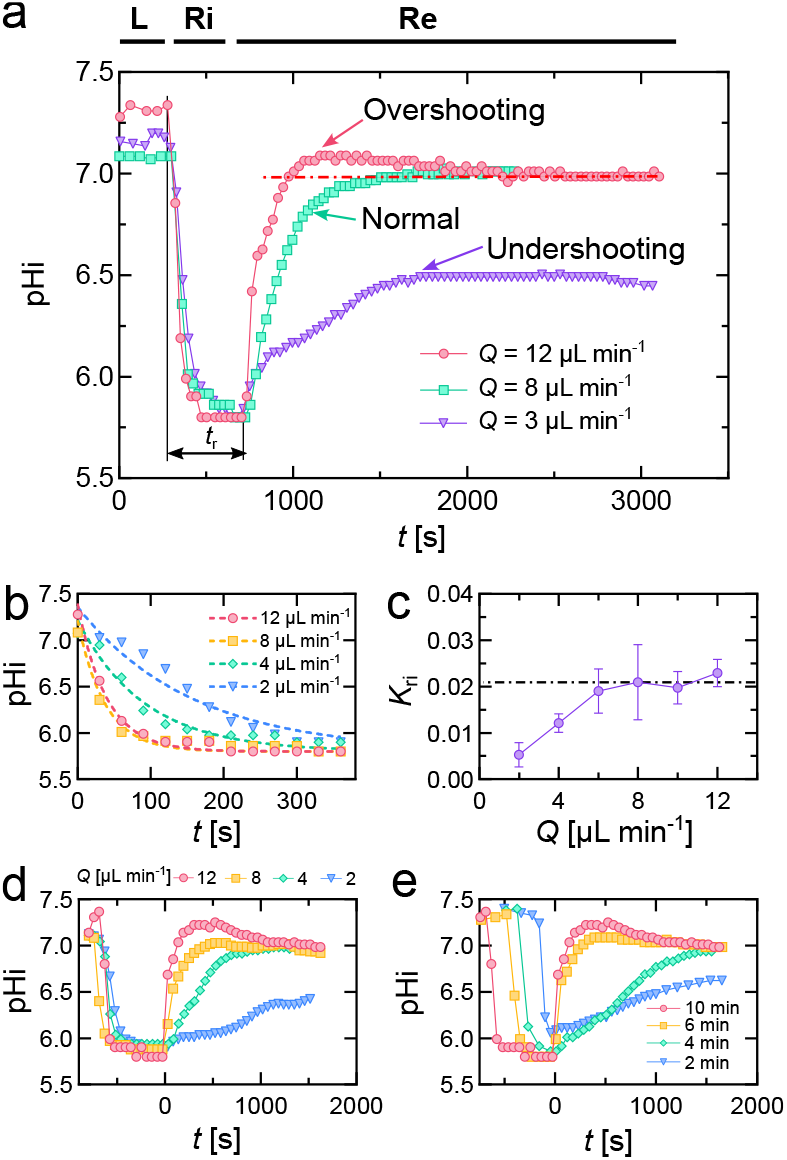
Behavior of intracellular pH during pH regulation assays. (a) Intracellular pH of representative cells during Loading, Rinsing and Recovery processes with rinsing duration *t*_r_ was kept at 6 min and *Q* varied from 3 to 12 µLmin^*−*1^. Over-shooting phenomenon happened at *Q* = 12 µLmin^*−*1^, while normal recovery happened at 8 µLmin^*−*1^, and 3 µLmin^*−*1^ resulted in undershooting. Dash-dotted line shows the intended pH value at steady state in the Overshooting phenomenon. (b) Representative data showing the decay of pH_i_ over time during Rinsing process at different flow rates. Dashed lines are exponential fits following Eq. 4. (c) The relation between the rinsing reaction rate *K*_ri_ and *Q*. Each data point represents the average value from *N* = 3 independent repeats (∼ 20 to 90 cells per condition per repeat). Error bars are standard deviations of the means. (d) Recovery of the intracellular pH of representative cells with rinsing duration *t*_r_ = 10 min and *Q* varied from 2 to 12 µLmin^*−*1^. (e) The effect of rinsing duration *t*_r_ on intracellular pH recovery while *Q* is fixed at 12 µLmin^*−*1^.

### 3.4 Influence of flow on intracellular pH during rinsing and recovery processes

As the pH_i_ dynamics were changed according to different flow rates, we investigated how the flow rate affected the kinetics of pH_i_ during rinsing and recovery process separately. For the Rinsing process, H^+^ is released in the cell as NH_3_ escapes through the cell membrane (Fig. 1(b)). By applying different flow rates during the Rinsing, we observed the pH_i_ decrease to a plateau at different rates (Fig. 3(b)). The reaction rate *K*_ri_ was obtained by fitting the decay of pH value with a single exponential, following the equation

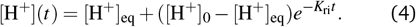

We achieved an increase of *K*_ri_ with higher rinsing flow rates, but the rate became saturated as *Q*≥ 6 µLmin^*−*1^ (Fig. 3(c)), with the value at saturation *K*_ri_ ≈ 0.021 ± 0.001s^*−*1^ (mean ± SD, *N* = 3 repeats). Since the rate of H^+^ released in the cell depends on the rate of NH_3_ diffusing through the cell membrane, we believe that the system reached a diffusion limit of NH_3_ when the flow rate is high enough.

On the other hand, we applied different flow rates during the Recovery process while keeping the rinsing duration at 10 min and monitored pH_i_ recovery (Fig. 3(d)). The long duration of the rinsing was applied to ensure that the rinsing had reached a steady state before the recovery process. Here, we observed the undershooting occurring at low flow rate (2 µLmin^*−*1^). At higher flow rates, pH_i_ turned to the normal recovery and the over-shooting state (Fig. 3(d)). We showed that flow rate plays an important role in the recovery dynamics of pH_i_. In fact, protons extruded by NHE1 result in a local increase in the concentration of proton outside of the cells, while Na^+^ local concentration outside of the cells decreases. ^22^ This leads to competitive inhibition, as protons can compete with Na^+^ for binding to NHE1, which lowers the exchange activity of NHE1. ^22,23^ Thus, fast flow rates in our microfluidic system help prevent this competitive inhibition and exhibit overshooting behavior of pH_i_.

Variation of flow rates is crucial on pH_i_ recovery because increasing flow rate reduces the thickness of the concentration boundary layer. When flow rate is sufficiently high, solutes transported by membrane proteins are quickly removed from the medium, preventing them from collapsing their own transmembrane gradients, which enables various behaviors and characteristic time scales for pH increase to be observed. We estimate the characteristic time scale for ions to move out of the boundary layer thickness of a cell with 5-micron length is around 0.03 seconds, ^24^ indicating that ion leakage is not limited in microfluidic flow. It also explains fast responses are not observed in our traditional perfusion experiments. ^13^ Moreover, even at high flow rate of 12 µLmin^*−*1^, the maximal shear stress acting on cells is approximately 0.16 Pa (1.6 dyne cm^*−*2^), which results in no effects on cells. ^25,26^

### 3.5 Rinsing of NH_3_ plays a big impact on pH recovery process

To elucidate how rinsing duration affects the recovery of pH_i_, we applied a constant flow rate of *Q* = 12 µLmin^*−*1^ with different rinsing duration, from 2 to 10 minutes. We obtained different pH_i_ recovery behaviors according to different values of *t*_r_. At *t*_r_ = 2 min, when the rinsing process did not reach the steady state yet, the pH_i_ recovery experienced a behavior similar to the undershooting state. When *t*_r_ = 4 min, the pH_*i*_ performed normal recovery. At higher rinsing duration (*t*_r_ ≥ 6 min), the pH_i_ recovered with overshooting behavior at high flow rate *Q* = 12 µLmin^*−*1^. We notice that when the rinsing process is not complete, NH_3_ remaining in cells can pick up protons to form 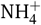. As a result, it affects intracellular proton concentration, which might slow down the recovery dynamics. In addition, we find that the duration of the rinsing should not be longer than 10 minutes, as cells that stay in acidic condition for a long time will become fatal.

### 3.6 State diagram of intracellular pH recovery

We performed a series of experiments with *t*_r_ ranging from 2 to 10 min, and *Q* ranging from 2 to 12 µLmin^*−*1^ to map a state diagram for pH_i_ recovery (Fig. 4). Here, we applied the same flow rate for the rinsing and recovery processes. We observed that low rinsing duration or low flow rates have the same effect on pH_i_ recovery, which resulted in undershooting state. High flow rates and enough rinsing duration helped the pH_i_ recover slowly or rapidly turn to an excited state with overshooting behavior. During the Rinsing process, the applied flow rate represents how fast NH_3_ dissipates after diffusing through the cell membrane and together with the rinsing duration determine the amount of residual 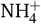 left inside the cell or the amount of intracellular 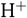 generated. Rinsing duration also reflects how long cells were put in acute condition. In the Recovery process, flow rate influences the competitive inhibition between the newly extruded H^+^ and the extracellular Na^+^, impacting the pH recovery speed. The state diagram shows that incomplete rinsing (residual intracellular 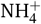) leads to an undershooting state, as more H^+^ is generated by 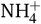 dissociation in competition with its extrusion by NHE1. On the other hand, complete rinsing with no competitive inhibition, especially when cells were under critically acute condition, results in an overshooting state. This further demonstrates that pH_i_ overshooting acts as a protective mechanism, allowing cells to adapt to and survive varying external conditions.

**Fig. 4.**
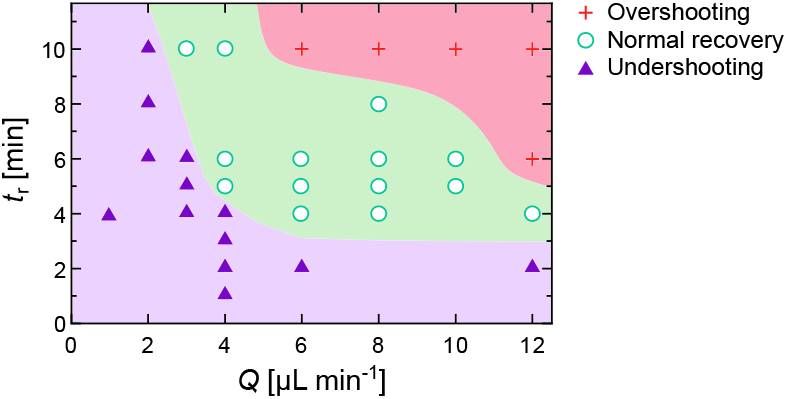
State diagram of the intracellular pH recovery under different flow rates *Q* and rinsing duration *t*_r_.

## Conclusion

In this study, we have developed a rapid microfluidic perfusion system for intracellular pH regulation assays on cells cultured on-chip. The system featured precise flow rate control of different solutions corresponding to different processes of pH changes and regulation (Loading, Rinsing, Recovery processes) across the cell region, together with live imaging by fluorescence microscopy and real-time data acquisition. We used pH-sensitive fluorescence dye BCECF/AM with double excitations for cell staining to probe pH_i_ changes over time. We also established a calculation tool to translate the ratios of fluorescence intensity emitted from the two wavelengths of the dye into the pH values in cells. By varying the flow rates of pH regulation solutions and the rinsing durations, we were able to induce a dynamic transition of the intracellular pH, from normal recovery to an overshooting or undershooting state. We also unraveled that the speed of intracellular pH change during the Rinsing process is capped at a limit despite increasing flow rate, which led to our observation that the speed of NH_3_ diffusing through the cell membrane is restricted. On the other hand, the recovery speed of intracellular pH, which is governed by the Na^+^/H^+^ exchanger NHE1, increases as the flow rate increases. If the flow rate is too low, the protons extruded by NHE1 will competitively inhibit NHE1 activity and in the meantime, local Na^+^ extracellular concentration will decrease. The fast flow rates allowed by our microfluidic device overcome such inhibitory effects and reveal overshooting that is challenging to produce otherwise.

The innovation of our work lies in the design of the microfluidic system that features rapid switching between solutions, fast and precise flow rate control, and realtime monitoring of intracellular pH. Such benefits enable first-time quantifiable and reproducible measurement of the overshooting of intracellular pH experimentally, a phenomenon that was not able to be achieved in this manner by our traditional perfusion system. ^13^ Future research will further characterize the pH overshooting behaviors and investigate the correlation between the level of pH overshooting and the experimental parameters (flow rate and rinsing duration, composition of solutions, combination of transporters activities). Altogether, we provided a robust platform for studying intracellular pH regulation, which can be further developed to monitor the intracellular pH in real time of complex cellular activities, such as cancer cells during migration or cells exposed to strong gradients and fast changes conditions.

## Author contributions

**Quang D. Tran**: Writing – original draft, Visualization, Validation, Methodology, Investigation, Formal analysis, Conceptualization. **Yann Bouret**: Writing – review & editing, Validation, Project administration, Methodology, Investigation, Formal analysis, Funding acquisition. **Xavier Noblin**: Validation, Supervision, Project administration, Methodology, Investigation, Funding acquisition. **Gisèle Jarretou**: Validation, Methodology, Investigation. **Laurent Counillon**: Validation, Supervision, Project administration, Methodology, Investigation, Funding acquisition. **Mallorie Poët**: Writing – original draft, Validation, Supervision, Project administration, Methodology, Investigation, Funding acquisition, Conceptualization. **Céline Cohen**: Writing – review & editing, Validation, Supervision, Project administration, Methodology, Investigation, Funding acquisition, Formal analysis, Conceptualization.

## Data availability

Data for this article are available at Zenodo at https://doi.org/10.5281/zenodo.13948923.

## Acknowledgements

We are thankful for the financial support from IDEX UCA JEDI (ANR-15-IDEX-0001, Nice, France): both from Academie IV (Project MpH) and Structuring Program BOOST (Project COMOZOO), Cancéropole PACA and Université Côte d’Azur. We thank Carina Krahé and Tom Montagnon for their preliminary experiments. We thank Michael Stehnach for fruitful discussions.

