## Supplementary Materials for "Rapid microfluidic perfusion system enables controlling dynamics of intracellular pH regulated by Na^+^/H^+^ exchanger NHE1"

### Translation of pH values from fluorescence ratios

**Fluorescence Intensity.** During the fluorimetry experiments, we use BCECF as $\alpha$ and its deprotonated form as $\beta$, with a $pK_a$ of 6.97 and the corresponding dissociation constant $K_a$:

$$\alpha \rightleftharpoons \beta + H^+, \;\; K_a = 10^{-pK_a} \approx 1.07 \cdot 10^{-7} = \frac{[\beta][H^+]}{[\alpha]}. \tag{1}$$

For a locally conserved concentration $C_0 = [\alpha] + [\beta]$ at a given $pH = -\log[H^+]$, and using $h = [H^+]$, we obtain the individual concentrations:

$$[\alpha] = C_0 \frac{h}{h + K_a}, \;\; \beta = C_0 \frac{K_a}{h + K_a}. \tag{2}$$

We define the respective fluorescence intensities of $\alpha$ and $\beta$ per unit of volume and per unit of concentration, and for a given illumination wavelength $\lambda$, as $\phi_\alpha(\lambda)$ and $\phi_\beta(\lambda)$. Accordingly, an observed volume $V$ emits the fluorescence intensity $I$ following:

$$\begin{aligned} I(h, \lambda, C_0) &= (\phi_\alpha(\lambda)[\alpha] + \phi_\beta(\lambda)[\beta])\, V \\ &= \left(\phi_\alpha(\lambda)\frac{h}{h + K_a} + \phi_\beta(\lambda)\frac{K_a}{h + K_a}\right) V C_0 \end{aligned} \tag{3}$$

**Isobestic point.** Let us define $\lambda_{iso}$ the isobestic wavelength such that both fluorescence intensities are equal to $\phi_{iso}$:

$$\phi_\alpha(\lambda_{iso}) = \phi_\beta(\lambda_{iso}) = \phi_{iso}. \tag{4}$$

For this particular wavelength we get:

$$I(h, \lambda_{iso}, C_0) = \phi_{iso} V C_0, \tag{5}$$

that is independent of $h$ and is used as a normalisation factor. For BCECF, $\lambda_{iso} \simeq 439$nm.

*
†
‡

**Intensity Ratio.** Let us use a working wavelength $\lambda_0$, and the define the *(reduced) fluorescence* as:

$$F(h) = \frac{I(h, \lambda_0, C_0)}{I(h, \lambda_{iso}, C_0)}. \tag{6}$$

We express $F(h)$ with two *fluorescence factors*:

$$\begin{cases} \Phi_\alpha = \dfrac{\phi_\alpha(\lambda_0)}{\phi_{iso}}, \\ \Phi_\beta = \dfrac{\phi_\beta(\lambda_0)}{\phi_{iso}}, \end{cases} \tag{7}$$

so that we rewrite:

$$F(h) = \frac{h\Phi_\alpha + K_a\Phi_\beta}{h + K_a}, \tag{8}$$

Hence we obtain higher sensitivity with respect to $h$ for distant values of $\Phi_\alpha$ and $\Phi_\beta$. this is achieved by choosing $\lambda_0 \simeq 490$nm near the maximum emission of the base form of BCECF (a.k.a $\beta$).

**Calibration.** We need to compute the effective values of $\Phi_\alpha$ and $\Phi_\beta$ since their value depends on the final optical setup. For that purpose, we perform two controlled measures the fluorescence, firstly in acidic conditions ($h = h_a$), then in basic conditions ($h = h_b$). We deduce the following equalities:

$$\begin{cases} F(h_a) = F_a = \dfrac{h_a\Phi_\alpha + K_a\Phi_\beta}{h_a + K_a} \\ F(h_b) = F_b = \dfrac{h_b\Phi_\alpha + K_a\Phi_\beta}{h_b + K_a}. \end{cases} \tag{9}$$

We solve the system above to get the fluorescence factors:

$$\begin{cases} \Phi_\alpha = \dfrac{1}{h_a - h_b}\left[F_a(K_a + h_a) - F_b(K_a + h_b)\right] \neq F_a \\ \Phi_\beta = \dfrac{1}{K_a(h_a - h_b)}\left[h_aF_b(K_a + h_b) - h_bF_a(K_a + h_a)\right] \neq F_b. \end{cases} \tag{10}$$

**pH Expression** We deduce the proton concentration as:

$$h = K\left[\frac{\Phi_\beta - F}{F - \Phi_\alpha}\right]. \tag{11}$$

Accordingly, we express the local pH as a function of the measured fluorescence:

$$\mathrm{pH} = \mathrm{pK} + \log\left[\frac{F - \Phi_\alpha}{\Phi_\beta - F}\right]. \tag{12}$$

Using this expression, the pH is evaluated for any fluorescence between $\Phi_a$ and $\Phi_b$ without any singularity, as ensured by the expression of $F$ in (8).
